# Identification and Utilization of Copy Number Information for Correcting Hi-C Contact Map of Cancer Cell Line

**DOI:** 10.1101/798710

**Authors:** Ahmed Ibrahim Samir Khalil, Siti Rawaidah Mohammad Muzaki, Anupam Chattopadhyay, Amartya Sanyal

## Abstract

**Motivation:** Hi-C and its variant techniques have been developed to capture the spatial organization of chromatin. Normalization of Hi-C contact maps is essential for accurate modeling and interpretation of genome-wide chromatin conformation. Most Hi-C correction methods are originally developed for normal cell lines and mainly target systematic biases. In contrast, cancer genomes carry multi-level copy number variations (CNVs). Copy number influences interaction frequency between genomic loci. Therefore, CNV-driven bias needs to be corrected for generating euploid-equivalent chromatin contact maps.

**Results:** We developed HiCNAtra framework that extracts read depth (RD) signal from Hi-C or 3C-seq reads to generate the high-resolution CNV profile and use this information to correct the contact map. We proposed the “entire restriction fragment” counting for better estimation of the RD signal and generation of CNV profiles. HiCNAtra integrates CNV information along with other systematic biases for explicitly correcting the interaction matrix using Poisson regression model. We demonstrated that RD estimation of HiCNAtra recapitulates the whole-genome sequencing (WGS)-derived coverage signal of the same cell line. Benchmarking against OneD method (only explicit method to target CNV bias) showed that HiCNAtra fared better in eliminating the impact of CNV on the contact maps.

**Availability and implementation:** HiCNAtra is an open source software implemented in MATLAB and is available at https://github.com/AISKhalil/HiCNAtra.

## 1. Introduction

Chromosome conformation capture (3C)-related methods are a collection of molecular techniques to analyze the three-dimensional (3D) chromatin interactions inside the cell [1]. In the last 15 years, the conventional 3C method [2] has been modified and combined with next-generation sequencing (NGS) to interrogate the chromatin interactions of genomic loci at different length scales [1, 3, 4]. In 2009, Hi-C protocol was developed to study the genome-wide chromatin interactions [3]. Hi-C experiments confirmed the chromosome territory hypothesis and established that our genome is hierarchically organized into A/B compartments and topologically-associating domains (TADs) [3, 5]. In addition, 3C-seq technique has been developed in recent years as a simple and straight-forward experimental protocol to study the genome-wide interactions of genomic loci. In this approach, a conventional 3C library is prepared first and then the library is sonicated and subsequently sequenced using NGS platform [4, 6, 7].

Majority of available Hi-C datasets were generated using cancer cell lines. However, cancer genomes are afflicted with large-scale and focal CNVs [8–12] that have pathological consequences [13]. Regions with copy number gains or losses will show concomitant changes in their interaction profiles [14–16]. This results in the rewiring of chromatin connectivity that may lead to alterations in the long-range control of gene expression [16]. CNVs can also modulate gene-regulatory mechanisms by altering the copy number of regulatory elements or by modifying the higher-order chromatin structure [17]. Study of prostate cancer cell lines showed that they have smaller-sized TADs with new boundaries. These new TAD boundaries occur at CNVs [18]. Recently, Wu *et. al.* reported in multiple myeloma (MM) cells with whole/partial chromosomal gains/losses that copy number (CN)-amplified regions have higher interaction frequencies than CN-neutral (normal) regions [15]. They identified the TADs by insulation score method [6] using ICE-normalized interaction matrices [19] and found that the numbers of TADs increased by 25% and the average size of TADs decreases significantly compared to the normal B cells. This suggests that CNV is an important source of bias on chromatin contact maps of cancer cell lines. Therefore, detection and correction of CNV effects are necessary to produce euploid-equivalent contact maps of cancer cells.

Hi-C sequencing reads can be utilized to estimate the read depth (RD) signal and discovery of CNVs without additional cost of performing WGS. Recently, HiCnv tool has attempted to identify CNVs from Hi-C reads [20]. However, it can only identify large CNVs (> 1Mb) which leave a lot of room for improvement in identifying smaller CNVs.

Correction of the raw contact frequencies is an essential step before any downstream analysis of the chromatin interaction data. Many normalization procedures have been developed over the years to correct the effects of these biases. For example, implicit methods usually model the Hi-C correction as matrix-balancing problem assuming that genomic regions of equal size should have similar coverage [19, 21–24]. On the other hand, explicit methods normalize the contact matrix by modelling the relationship between the contact frequencies and the inherent systematic biases introduced by GC content, restriction fragment length and mappability biases only [25, 26].

Though implicit method may indirectly normalize the CNV-associated biases of the interaction matrix, however, this method relies on empirical conditions to normalize the contact frequencies without a sound biological basis. For example, ICE method [19] empirically removes the top 0.5% of most frequently-detected restriction fragments as well as 1% of bins with lowest read abundance (read/bin filtering steps). Also, the iterative correction step is performed till the variance of the bias sources becomes negligible which is again dependent on the magnitude of the biases [19]. These filtering steps and thresholds may lead to matrix normalization to a reasonable level for normal cell lines. However, they cannot be generalized for all cancer cell lines where the contact frequencies are influenced significantly by CNVs [14–16].

Recently, OneD [27] tool has been developed for explicitly correcting the CNV bias from Hi-C contact maps. Evaluation of OneD versus other explicit and implicit Hi-C correction methods shows that OneD outperforms them for correcting Hi-C contact maps of cancer cell lines [27]. OneD normalizes the contact map sequentially for the sample-independent systematic biases (such as GC content, restriction fragment length and mappability) followed by sample-driven CNV bias. First, a generalized additive model (GAM), based on the negative binomial (NB) distribution, is utilized for normalizing contact frequencies from the sample-independent biases. Then, OneD divides the contact frequencies between two genomic loci by the product of the copy numbers (estimated by a hidden Markov model) of these loci. However, in case of CNV-infested cancer cell lines, GAM cannot estimate the accurate contribution of each systematic bias on the contact map without including CNV bias with other biases at the same time. Therefore, we believe that OneD method may not normalize the contact frequencies of cancer cell lines optimally.

Here, we present HiCNAtra framework that computes the RD signal from the NGS reads of Hi-C experiment to construct the high-resolution CNV profile and use this profile as one of the bias sources to normalize the contact matrix. HiCNAtra utilizes all the raw Hi-C reads using a novel “entire restriction fragment” counting approach. This enables estimation of RD signal at high resolution that allows precise detection of both large-scale and focal alterations based on CNAtra approach [28]. For 3C-seq, we generate the RD signal and CNV profiles exclusively from the genomic reads. We then utilize a generalized linear model (GLM) to correct the interaction matrix from the sample-driven CNV bias along with sample-independent systematic biases. HiCNAtra also provides an interactive platform to visualize and manually inspect the complete CNV profile, contact map and accessory information for further validation and interpretation. We benchmarked the performance of HiCNAtra against OneD [27] using Hi-C/3C-seq data of five cancer cell lines. Our RD-computing approach from Hi-C/3C-seq data from cancer cell lines successfully recapitulate the RD signal derived from WGS data of the corresponding cell lines. Moreover, HiCNAtra-normalized contact maps are least correlated with all sources of biases. Manual review and visualization also validated the advantage of HiCNAtra over OneD method in ameliorating the effects of CNV-induced artifacts on contact maps.

## 2. Methods

The detailed description of methods is provided in Extended Methods under Supplementary Information. Here, we briefly explain the HiCNAtra framework and NGS data processing. The HiCNAtra software is freely available along with the user manual from https://github.com/AISKhalil/HiCNAtra. The user manual contains all the necessary information about the input data and all output results.

### 2.1 HiCNAtra framework

HiCNAtra pipeline is divided into three modules (Fig. 1a): 1) computation of the RD signal from Hi-C or 3C-seq reads (*RD calculator*), 2) RD-based detection of copy number events (*CNV caller*) and 3) bias correction of interaction matrix introduced by CNVs and other systematic biases (*Contact map normalization*). Briefly, HiCNAtra pipeline starts with computation of RD signal from the Hi-C/3C-seq reads. Next, we use CNAtra [28] approach to identify large-scale and focal alterations from the RD signal and integrate them to generate the CNV track. This CNV track is used as an explicit bias source along with other systematic biases for correcting interaction matrix. Finally, we utilize a Poisson-based GLM to normalize the contact map for GC content, mappability, and fragment length biases as well as biases introduced by copy number gains/losses.

**Fig. 1.**
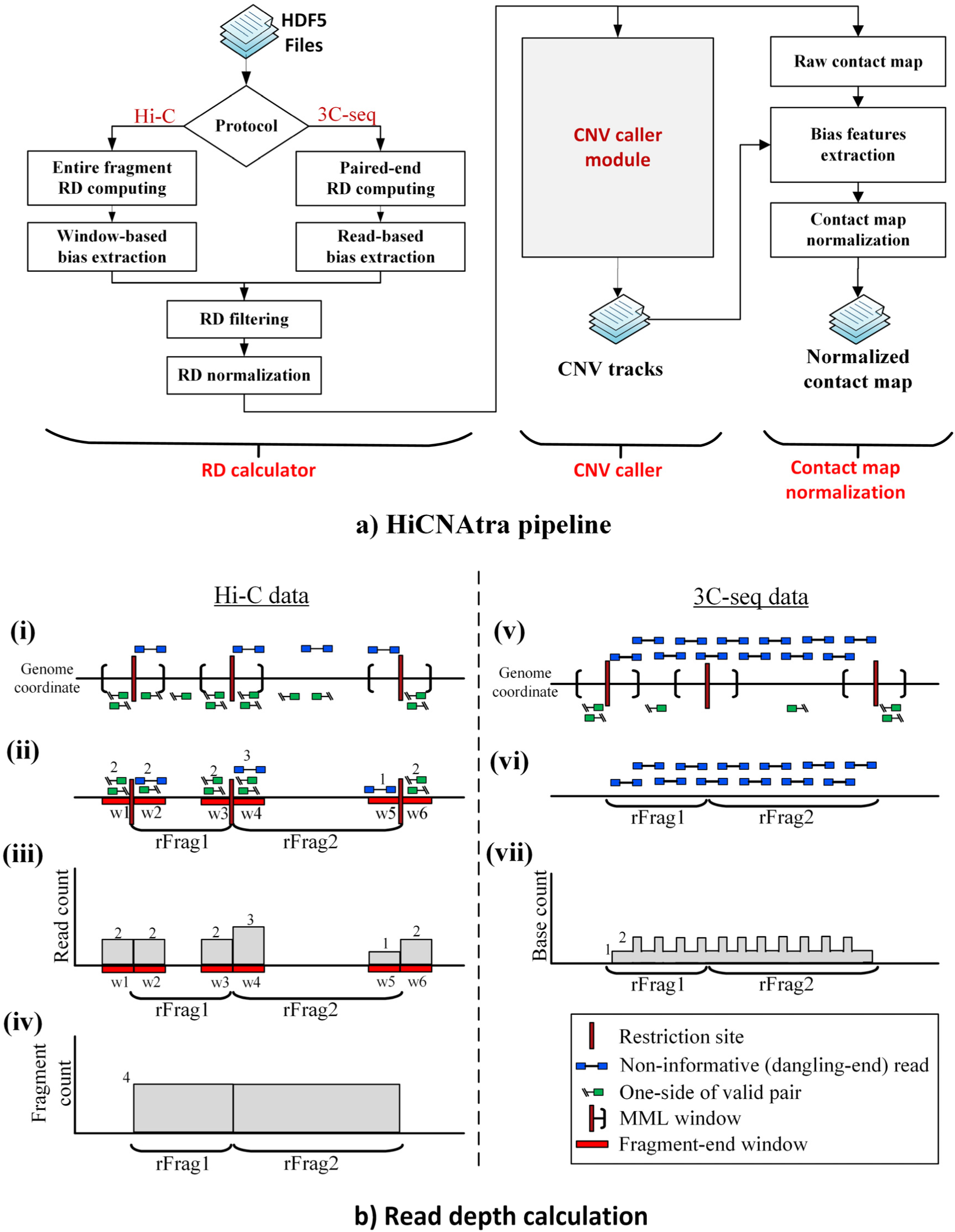
HiCNAtra methodology and RD signal computation. **(a)** Block diagram of HiCNAtra software. The HiCNAtra pipeline is divided into RD calculator, CNV caller, and contact-map normalization modules. **(b)** Cartoon illustrations of the RD signal computation method from Hi-C (left) and 3C-seq (right) data. For Hi-C, informative and non-informative (only dangling-end shown) reads **(i)** that were mapped within the restriction fragment-end windows **(ii)** are used for RD calculation. Then, number of reads is counted for each window **(iii)**. Base counts are then estimated for each restriction fragment as the summation of the counts of its two fragment-end windows **(iv)**. For 3C-seq libraries **(v)**, genomic reads **(vi)** are exclusively used to compute the RD signal in an unbiased manner like WGS paired-end reads **(vii)**.

#### 2.1.1 Computing the RD signal from Hi-C/3C-seq NGS reads

Hi-C/3C-seq datasets comprise different read types such as valid pairs, dangling-end, extra dangling-end, self-circle, and single-sided reads as well as genomic reads [29]. These reads can be mainly categorized into (1) ‘informative’ reads containing valid pairs that represent interactions between genomic loci, and (2) ‘non-informative’ reads that include all other types of reads. In case of Hi-C, valid pairs generally comprise ~40-65% of the total mapped reads and these reads are solely used to generate contact map. In addition to valid pairs, HiCNAtra also utilizes the non-informative reads to compute the RD signal at higher resolution. On the other hand, in the absence of biotin labeling and pull-down steps, the majority of 3C-seq reads are contributed by genomic reads (~70-80% of the double-sided mapped reads). Therefore, for 3C-seq datasets, we compute the RD signal exclusively from these genomic reads in a similar manner as the RD signal is usually computed from paired-end WGS reads.

Based on the Hi-C or 3C-seq experiment, reads are utilized differently for computing the RD signal (default bin size = 5 Kb). For Hi-C datasets, HiCNAtra computes the RD signal from all Hi-C reads (Fig 1b(i)) using the “entire restriction fragment” counting approach. For this, we first retain only those reads that are located within the restriction fragment-end windows-window of maximum molecule length (MML) next to the restriction site (Fig 1b(ii)). The reason for targeting fragment-end window reads is that the biotin pull-down step of Hi-C protocol should cause most of the reads to map next to the restriction sites. Second, we count the reads for each restriction fragment based on the assumption that each continuous DNA sequence read represents a particular restriction fragment and contribute to the abundance of that fragment. Therefore, we count each non-informative read as a single count whereas for valid pairs, side1 and side2 are counted separately (Fig 1b(iii)). So for a particular restriction fragment, the fragment count is calculated as the sum of the number of reads located in both fragment-end windows (Fig. 1b(iv)). For example, the fragment count of restriction fragment 1 (rFrag1) is the sum of read counts of fragment-end windows, w2 and w3 (Fig. 1b(iv)). Finally, we assign the fragment count of a restriction fragment to all of its bases and compute the RD signal. For 3C-seq datasets, genomic reads are distributed uniformly along the genome and represent the majority of reads (Fig 1.1b(v)). Therefore, we use these paired-end genomic reads (Fig 1.1b(vi)) to compute the RD signal in an unbiased manner (Fig. 1.1b(vii)).

#### 2.1.2 CNV identification

The CNV calling module of HiCNAtra is based on CNAtra [28] approach. CNAtra CNV caller constitutes the hierarchical framework to delineate the multi-level copy number alterations in the cancer genomes. Briefly, we first define the CN reference (CN=2) by fitting the RD signal to a multimodal distribution. Then we utilize a multi-step framework to first identify large genomic segments with distinct CN state. Segments with CN state other than 2 (CN≠ 2) are considered as large-scale copy number variations (LCVs) or segmental aneuploidies. Next, focal amplifications and deletions in each CN-defined segment are detected based on coverage-based thresholding. HiCNAtra then computes the CNV tracks by merging both LCVs and focal alterations (FAs) for Hi-C correction.

#### 2.1.3 Correction of the chromatin contact map

We employed GLMs to normalize the contact frequencies against all sources of biases similar to HiCNorm [26]. Two GLM with Poisson distribution are used to fit the *cis* and *trans* contact maps separately. By default, we apply our normalization approach in a genome-wide manner for better estimation of the GLM parameters.

##### Normalization of *cis* contact map

Let 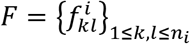 represent the *n*_*i*_ × *n*_*i*_ *cis* interaction frequencies for chromosome *i*, where *n*_*i*_ is the number of bins in chromosome *i*. Each element 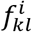 represent the number of interactions between genome loci from bin *k* and bin *l* in chromosome *i*. Let 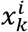, 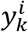, 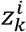, and 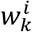 represent the GC content, mappability, effective length, CNV features of bin *k* in chromosome *i*, respectively; whereas 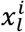, 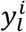, 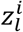, and 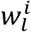 represent the features of bin *l* in chromosome *i*, respectively. We assume that the interaction frequency 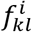 follows Poisson distribution with rate *λ*_*jk*_:

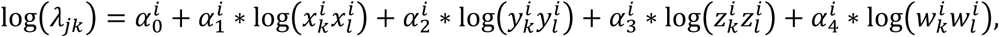

where *α*^*i*^_0-4_ are the coefficients of the GLM model. We fit this GLM model and use the coefficient estimates 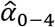 for computing the estimated Poisson rates:

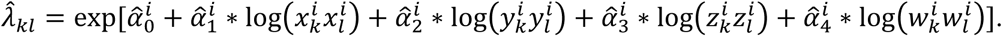

Then, 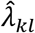 is used to compute the normalized interaction frequency 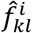 between bin *k* and bin *l* in chromosome *i* of bin as following:

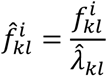

##### Normalization of *trans* contact map

Let 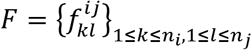 represent the *n*_*i*_ × *n*_*j*_ *trans* interaction frequencies between chromosome *i* and chromosome *j*, where *n*_*i*_ and *n*_*j*_ are the number of bins in chromosomes *i* and *j*, respectively. Each element 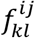 represent the number of interactions between genome loci of bins *k* and *l* from chromosomes *i* and *j*, respectively. Let 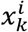, 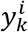, 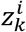, and 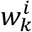 represent the GC content, mappability, effective length, CNV features of bin *k* in chromosome *i*, respectively; whereas 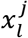, 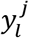, 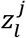, and 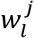 represent the features bin *l* in chromosome *j*, respectively. We assume that the interaction frequency 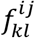 follows Poisson distribution with rate *λ*_*jk*_:

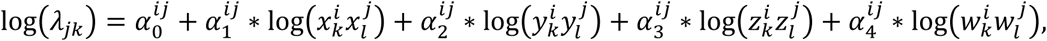

where *α*^*ij*^_0-4_ are the coefficients of the GLM. We fit this GLM model and use the coefficient estimates 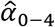 for computing the estimated Poisson rates:

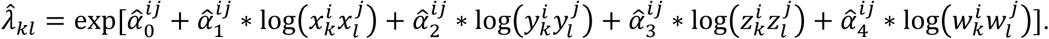

Then, 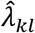 is used to compute the normalized interaction frequency 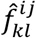 between bins *k* and *l* from chromosomes *i* and *j*, respectively:

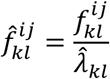

### 2.2 Data processing

We used six publicly available Hi-C datasets (GM12878, IMR90, PrEC, MCF7, LNCaP and PC3) and two in-house generated 3C-seq datasets (K562 and NCI-H69). We used input control sequencing reads of ChIP-seq experiments of MCF7, LNCaP and K562 cells to generate WGS-derived RD signal to validate HiCNAtra RD signal. Detailed information about the sequencing datasets is provided in the Supplementary Table 1. The wet lab experimental details about cell culture and 3C-seq library preparations are provided in Extended Methods under Supplementary Information.

For both Hi-C and 3C-seq datasets, we applied the iterative-mapping technique of hiclib [19] for aligning short sequence reads to human reference genome GRCh37 (hg19). The mapping output HDF5 files are used as input for HiCNAtra tool. Classification of read types (valid pairs, dangling-end, extra dangling-end, self-circle) depends on the maximum molecule length (MML). For each dataset, we approximately set the MML as the size (in hundreds of bp) that is greater than the fragment lengths (side1-start to side2-end) of 99% of dangling-end and extra-dangling end reads (Supplementary Table 1, Supplementary Fig. S1).

## 3. Results

### 3.1 Utilization of all Hi-C reads for computing high-resolution RD signal

Data coverage (number of reads) is an important factor for detecting the CNVs, especially the focal alterations. Although valid pairs are the main target of Hi-C/3C-seq protocols for capturing the chromatin interactions, other non-informative reads can be additionally utilized for estimation of the RD signal at a higher resolution. Based on the analyses of six Hi-C datasets, valid pairs represent 39-64% of the total mapped reads (Supplementary Table 1). Other reads, which represent a significant chunk (36-61%), are usually filtered out during Hi-C downstream analysis. However, these reads are still ‘*informative*’ and can potentially contribute to the estimation of the genome-wide RD signal. Therefore, we computed RD signal using both valid pairs and non-informative reads. Addition of these non-informative reads increases the resolution of the RD signal by 22-44% for different Hi-C datasets (Supplementary Table 1). The correctness of incorporating non-informative reads to estimate the RD signal is evaluated by comparing “read counts per restriction fragment” from all Hi-C reads versus valid pairs only. We found that read counts are highly correlated (Spearman correlation 80-91%) (Supplementary Table 2) confirming the validity of using non-informative Hi-C reads in RD calculation which results in higher resolution of RD signal. This conferred advantage to HiCNAtra over HiCnv and OneD which use only the valid pair reads for estimating the RD signal.

On the other hand, 3C-seq technique, which combines 3C library with NGS sequencing step, results in a higher percentage of genomic reads among the double-sided mapped reads (66% for K562 and 77% for NCI-H69). Therefore, similar to WGS datasets, RD signals can be conveniently computed from these genomic reads. Each paired-end genomic read is used to calculate the fragment length (side1-start to side2-end). This results in computing RD signal at high coverage (14.26x for K562 with only 117 million genomic reads and 12.37x for NCI-H69 with only 120 million reads). This demonstrates that genomic reads from the 3C-seq datasets can be effectively used for estimating the RD signal in an unbiased manner without additional cost of WGS.

### 3.2 Entire restriction fragment counting approach successfully extracts copy number-associated features of RD signal

Accurate estimation of the RD signal from Hi-C data is the primary key for identification of copy number events. Generally, most Hi-C analysis tools [19–27] assign the Hi-C reads to the midpoint of corresponding restriction fragment (midpoint approach). Alternatively, exact cut site of the read can be used (exact-cut approach) [30]. These approaches can be suitable for contact frequency calculation since Hi-C matrices are usually generated using large bin size. Usually, the bin size is at least one order of magnitude larger than the restriction fragment length. However, smaller bins are essential for effective identification of change points at high-resolution for discovery of both LCVs and FAs. Therefore, we introduced “entire restriction fragment” counting approach for RD calculation (Fig. 1b). This allowed us to compute the RD signal at high-resolution. We compared HiCNAtra RD calculation method with exact-cut and midpoint approaches using MCF7 Hi-C data generated using HindIII enzyme (Fig. 2a). We calculated the RD signal at 5-Kb bin using Hi-C reads which is of the same order of magnitude as the experimental resolution (average 4096 bases for 6-bp cutter). Interestingly, we found that our approach can effectively capture LCVs as evidenced by two peaks in the RD signal distribution (Fig 2a right, bottom panel). In addition, FAs are conspicuous using this approach viz. FA1 and FA2 (Figure. 2a left, bottom panel). In comparison, the RD signal generated using other approaches show overdispersion and make the distribution more skewed toward bins with low reads (Figure 2a, top and middle panels).

**Fig. 2.**
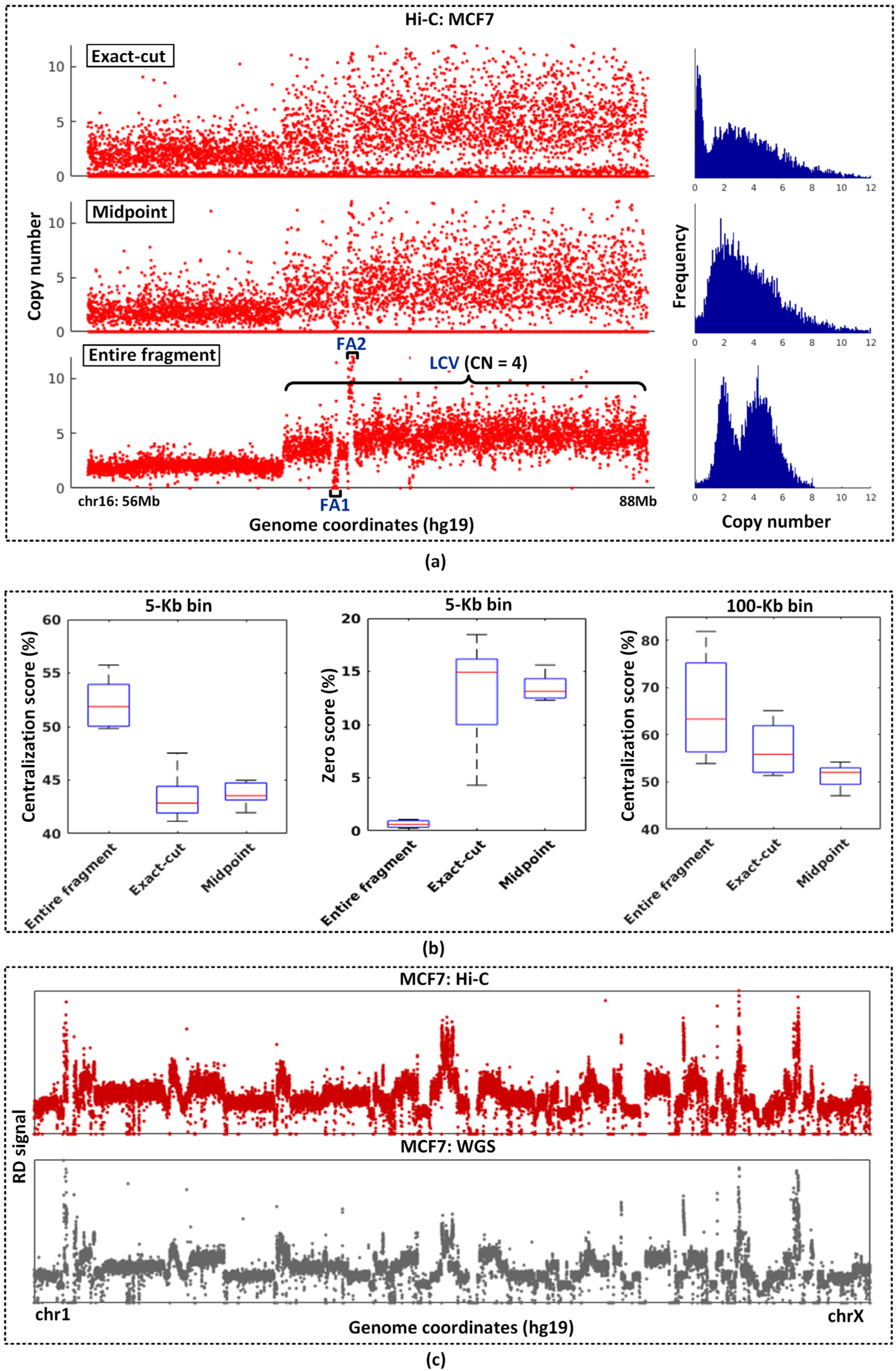
Entire restriction fragment approach is the best estimate of RD signal. **(a)** The coverage plot (left) at 5-Kb bin and the RD frequency distribution (right) of MCF7 Chr16:56-88Mb region extracted from Hi-C data using exact-cut (top), midpoint (middle), and HiCNAtra entire fragment (bottom) approaches. Each red dot represents copy number of bin. **(b)** Centralization score and zero score of the Hi-C RD signals using “entire restriction fragment” counting, exact-cut, and midpoint approaches using bin sizes of 5 Kb and 100 Kb. **(c)** The genome-wide coverage plot of MCF7 cancer cell computed from Hi-C data (top) and WGS data (bottom). WGS data is obtained from ChIP-seq input control sequencing reads. Each red dot is the RD value per bin from Hi-C data whereas each grey dot represents RD value per bin from the WGS data.

For a more quantitative comparison, we computed the centralization score (CS) and zero score (ZS) for all the six Hi-C datasets. CS is computed as the percentage of bins that are near their integer CN states. CS can be used as a measure for distributing genomic loci into distinct CN states. We then defined ZS as the percentage of bins with CN=0. ZS can be used as a measure of the sparseness of the RD signal. We found that “entire fragment” counting approach achieved the maximum CS (for both 5-Kb and 100-Kb bin) and minimum ZS in all Hi-C datasets (Fig. 2b, Supplementary Table 3). This is also confirmed by the visual inspection of coverage plot and RD frequency distribution derived from these three approaches (Fig. 2a). High CS implies that RD signal is distributed near distinct CN states which aids in the identification of CNVs. Low ZS is essential to avoid false detection of deleted regions, especially in large restriction fragments.

Next, we validated that the “entire restriction fragment” counting approach extract the RD signal correctly from Hi-C reads. We compared the RD signal derived from Hi-C datasets of MCF7 and LNCaP with the WGS-derived coverage signal of same cell lines. First, we found that HiCNAtra RD signal has the highest correlation with the WGS-derived RD signal for both MCF7 (Spearman’s ρ =0.714) and LNCaP (Spearman’s ρ =0.55) cells (Supplementary Table 3). In comparison, exact-cut and midpoint approaches have the Spearman’s ρ of 0.32 and 0.31 for MCF7, and 0.26 and 0.24 for LNCaP cells. This confirmed that the HiCNAtra counting approach is the best estimate of the RD signal. Similarly, we computed the RD signal (5-Kb bin) from genomic reads of K562 3C-seq dataset. As expected, RD signal extracted from K562 3C-seq data correlated well with WGS-derived coverage signal of K562 cells (Spearman’s ρ = 0.573). This confirmed that the RD signal can be reproducibly derived from Hi-C/3C-seq datasets (visually illustrated in Fig. 2c and Supplementary Fig. S2).

### 3.3 HiCNAtra can extract high-resolution copy number events from Hi-C/3C-seq datasets

Successful cloning of the WGS-derived RD signal from Hi-C/3C-seq reads enables HiCNAtra to accurately detect CNV regions. To the best of our knowledge, HiCnv [20] is the only tool available to detect CNV from Hi-C data. However, HiCnv computes the RD signal using midpoint approach exclusively from the valid pairs. As stated before, valid pairs represent 39-64% of total mapped-Hi-C reads. Therefore, HiCnv sacrifices a large percentage of reads which can be effectively used for calculating the RD signal based on HiCNAtra approach. Moreover, the midpoint approach limits the power of HiCnv to detect only large CNVs (size > 1Mb) [20]. As an alternative solution, we adapted the hierarchical CNV detection approach of CNAtra [28] in HiCNAtra CNV caller module to identify both LCVs and FAs.

We analyzed the CNV profiles of all six Hi-C and two 3C-seq datasets using HiCNAtra (Supplementary Table 4). Results show that all cancer cell lines (MCF7, LNCaP, PC3, K562, NCI-H69) are enriched for LCVs (41-122 regions) with a median width of 5.7-45.7 Mb, whereas normal cell lines (GM12878, IMR90 and PrEC) are almost free of LCVs (1-2 regions) (Supplementary Fig. S3, bottom panel). On the other hand, focal alterations are pervasive in both cancer (162-238 regions) and normal cell lines (25-107 regions) with median width ranging from 157 to 225 Kb (Supplementary Fig. S3, top panel). Focal alterations (FAs) are divided into focal amplifications (5-161 regions), homozygous deletions (CN=0) (10-72 regions), and hemizygous deletions (CN=1) (7-95 regions). Visual inspection of the CNV profiles generated from MCF7 Hi-C data and NCI-H69 3C-seq data clearly illustrated that chromosomes are interspersed with both large-scale and focal alteration events (Supplementary Fig. S4). Next, CNV track is generated by integrating LCVs and FAs. This track is used for correcting the contact frequencies of Hi-C/3C-seq datasets.

### 3.4 HiCNAtra successfully ameliorates the effects of CNVs on the contact map

It is well-established that chromatin contact map is highly influenced by the copy number events [15, 18]. Our analysis of Hi-C/3C-seq data demonstrated that interaction frequencies are positively correlated across cancer cell lines with the copy number tracks derived from the same cell line. This correlation is almost similar to effective length (restriction fragment) bias and greater than GC-content and mappability biases (Fig. 3a, Supplementary Fig. S5).

**Fig. 3.**
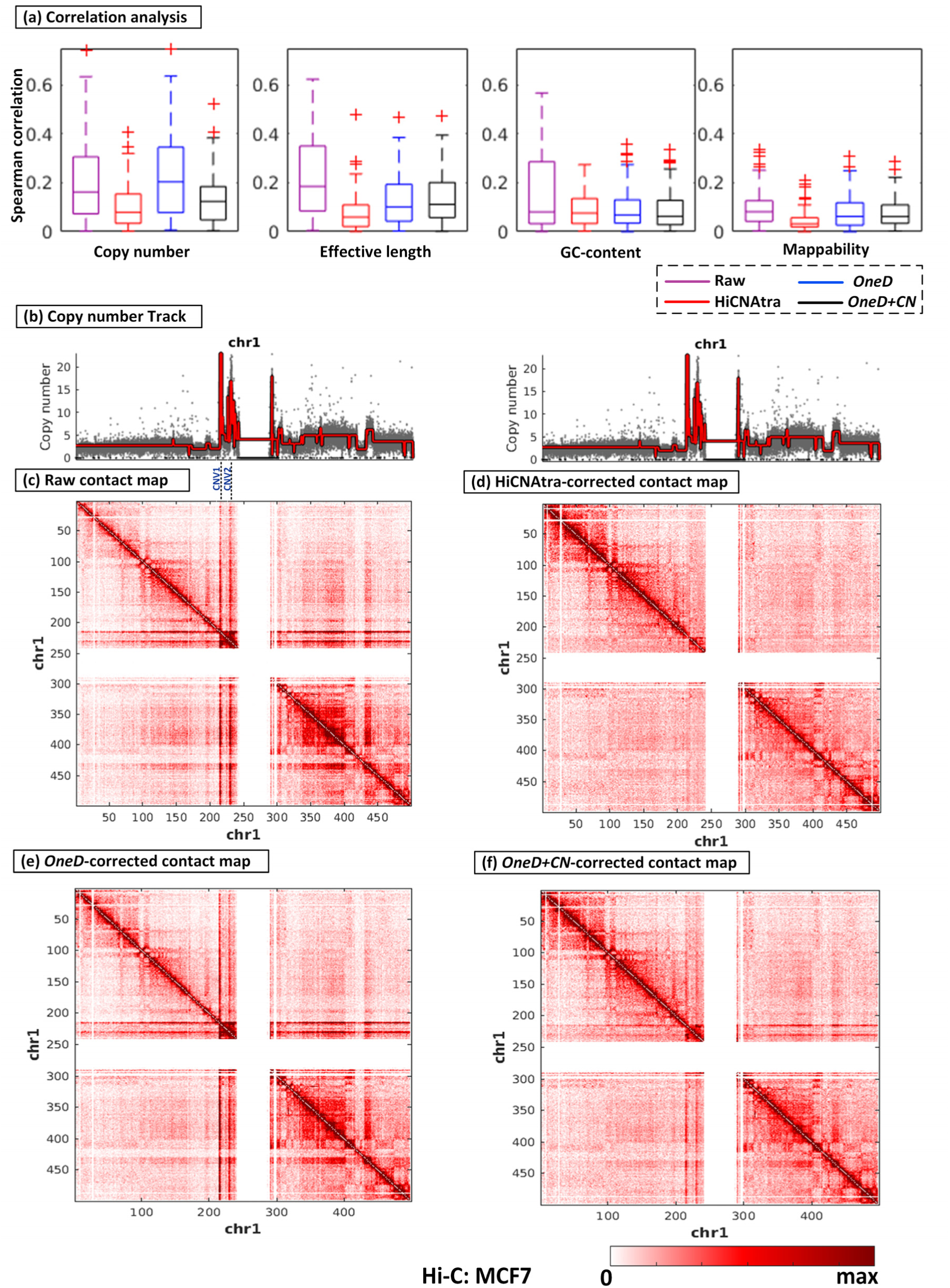
HiCNAtra normalization successfully ameliorated the effects of CNVs on the Hi-C contact maps of cancer cells. **(a)** Spearman correlations between *cis* contact frequencies of cancer datasets (MCF7, LNCaP, PC3, K562, NCI-H69) across chromosomes and sample-independent and dependent biases using HiCNAtra, *OneD*, *OneD+CN* approaches. **(b)** CNV track of chr1 of MCF7 Hi-C data. Each grey dot represents the copy number of a bin. The red line represents the copy number track where any amplitude transition indicates a new CNV region. **(c-f)** Pre- and post-normalized Hi-C interaction heatmaps (500-kb bin) of MCF7 chr 1. Raw contact map **(c)** and HiCNAtra-corrected **(d)**, *oneD*-corrected **(e)**, and *oneD+CN*-corrected **(f)** contact maps are shown. The color scale denotes interaction frequencies between bins. It can be visually observed that amplified and deleted regions (Fig. 3b) show a concomitant increase or decrease of contact frequencies in the raw Hi-C heatmap (Fig. 3c). For example, amplified regions CNV1 and CNV2 (Fig. 3b) resulted in two off-diagonal block patterns (visible artifacts) with signal higher than their surrounding regions (Fig. 3c).

Next, inspired by HiCNorm [26], we utilized GLM with Poisson distribution to normalize the contact frequencies (the variable) versus CNV, effective fragment length, mappability, and GC-content (the predictors). In order to evaluate the performance of HiCNAtra, we benchmarked it against OneD [27], the only available tool for explicitly correcting CNV effects on contact maps of cancer cells.

Computationally, OneD first utilizes a GAM with negative binomial (NB) distribution to correct the contact frequencies from the effective fragment length, mappability, and GC-content. Then, it normalizes the resulted interaction frequencies again for the CNV bias, estimated by a hidden Markov model. They divide the interaction frequency *f*_*ij*_ between two bins *i* and *j* by the copy number of these bins (*c*_*i*_, *c*_*j*_) as 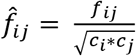. However, OneD’s sequential normalization may not normalize the biases of cancer cell lines optimally. Theoretically, when CNV is a dominant bias, GAM cannot accurately compute its coefficient estimates between the variable (interaction frequency) and three predictors (effective length, mappability, GC-content) since it does not include the main predictor (CNV bias). Therefore, it cannot capture the correct contribution of each systematic bias (effective length, mappability, GC-content) on the interaction frequencies. In contrast, HiCNAtra includes CNV bias with other systematic biases for *simultaneously* correcting the contact frequencies of Hi-C/3C-seq data. This allows HiCNAtra to learn the accurate relationship between the contact frequencies and their biases.

We have earlier demonstrated the advantage of HiCNAtra in computing high-resolution RD signal from Hi-C reads using the “entire restriction fragment” counting approach over midpoint approach, which is used by OneD. Therefore, for fair comparison, we implemented the OneD correction methods (*OneD* and *OneD+CN*) using HiCNAtra generated CNV track. The *OneD* module includes the GAM with NB distribution for correcting the effective length, mappability, and GC-content biases only. *OneD+CN*, on the other hand, includes the GAM for correcting the systematic biases, followed by the CNV bias correction. First, we normalized all the five Hi-C/3C-seq contact maps of cancer cell lines using three approaches (HiCNAtra, *OneD* and *OneD+CN*). Then, we computed the Spearman correlations between the biases and corrected contact frequencies. Overall, our results showed that HiCNAtra-corrected interaction frequencies achieved the minimum correlation with CNV tracks (median Spearman’s ρ = 0.08) as well as with other systematic biases across cell lines (Fig. 3a, Supplementary Fig. S5). The *OneD+CN*-corrected contact frequencies showed the second-lowest correlation with CNV tracks (mean Spearman’s ρ = 0.124) and systematic biases (Fig. 3a, Supplementary Fig. S5). This confirms the advantage of HiCNAtra’s *simultaneous* bias correction method over *OneD+CN*’s *sequential* bias correction.

Visualization and manual inspection of contact heatmaps is a simple way to qualitatively ascertain the effects of CNVs on interaction frequencies. Cancer cell lines, which typically harbor CNVs, create visible artifacts on the raw heatmaps. For example, the amplified and deleted regions in CNV track (Fig. 3b) of MCF7 chr1 show a concomitant increase or decrease of contact frequencies in the raw (uncorrected) heatmap (Fig. 3c). Specifically, amplified regions CNV1 and CNV2 (Fig. 3b) resulted in two off-diagonal block patterns (artifacts) in the heatmap with signal higher than their surrounding regions (Fig. 3c). These blocks refer to the overrepresented *cis* interactions between these CNV regions with other genomic regions. We plotted bias-corrected heatmaps of the MCF7 chr1 normalized using HiCNAtra, *OneD* and *OneD+CN* and compared them with the raw heatmap to visually interpret their abilities to ameliorate CNV-induced artifacts. Visually, it can be noticed that off-diagonal lines (contributed by CNV1 and CNV2) are removed in the HiCNAtra-normalized heatmap without distorting its overall features (Fig. 3d). As expected, *OneD-* normalized heatmap is almost similar to the raw heatmap in the absence of CNV bias correction (Fig. 3e). Although *OneD+CN*-normalized heatmap showed improvement over the raw heatmap, however, the footprint of off-diagonal blocks (artifacts) of weaker strength is still visible after bias correction (Fig. 3f). We witnessed similar observations while inspecting other regions of MCF7 as well as in a different cancer cell line (Supplementary Fig. S6). This confirms that CN information should be integrated with other systematic biases for effective normalization of Hi-C/3C-seq contact frequencies of cancer cells. In case of normal cells, which are largely devoid of CNVs, HiCNAtra approach of systematic bias correction of contact maps is basically same as HiCNorm approach. Therefore, HiCNAtra approach can be applied for both normal and cancer cell lines.

## 4. Discussion

Cancer genomes are scattered with large-scale CNVs and focal alterations. Amplification and deletion events result in over- and under-representation of chromatin interactions respectively in Hi-C/3C-seq datasets. Therefore, CNVs can greatly influence the interpretation of chromatin contact maps and may lead to false identification of genomic regions with over- or under-represented interaction strengths. The effect CNV bias should be compensated for the amplified/deleted regions along with other systematic biases for accurate interpretation of interaction frequency.

High-resolution extraction of RD signal from Hi-C/3C-seq datasets is central to discover copy number events of smaller sizes. We propose two solutions to better estimate RD signal from Hi-C datasets. First, HiCNAtra utilized all types of Hi-C reads that result in higher-resolution RD signal unlike other tools which use only valid pairs. Second, in order to better demarcate the RD signal into distinct copy number state, HiCNAtra employs the “entire restriction fragment” counting approach. Consequently, HiCNAtra can better extract the RD signal from the Hi-C reads, which corroborated well with the WGS-derived RD profile.

Previous studies have demonstrated the adverse effects of CNV regions on TAD sizes and structures in cancer cells. We observed a positive correlation between the copy number and the strength of the interaction frequencies of genomic loci. This clearly suggests that CNV imparts a dominant explicit bias in case of cancer cell lines. HiCNAtra identified the copy number events and incorporated them with other systematic biases for simultaneously correcting the contact map. Compared to OneD approach, HiCNAtra correction method is better suited to normalize the contact maps of cancer cell lines as evidenced by visual inspection of raw and corrected heatmaps.

In conclusion, our results suggest that HiCNAtra provides a better solution for a) computing high-coverage RD signal and detecting large-scale and focal CNVs from Hi-C/3C-seq datasets, and b) for explicitly correcting the chromatin contact frequencies from biases introduced by CNVs.

## Declarations

## Acknowledgment

We like to thank Ms. Yao Chen for help in analyzing 3C-seq datasets and fruitful discussions. We also acknowledge Sanyal and Chattopadhyay lab members for their valuable comments.

## Author’s contributions

AS, AC and AISK conceived the project. AISK developed HiCNAtra software with inputs from AS and AC and performed all the analyses. SRMM prepared the 3C-seq libraries. AS, AISK and AC analyzed the data and prepared the manuscript. All authors read and approved the final manuscript.

## Ethics approval and consent to participate

Not applicable.

